# A phenotypically robust model of Spinal and Bulbar Muscular Atrophy in *Drosophila*

**DOI:** 10.1101/2023.03.25.534140

**Authors:** Kristin Richardson, Medha Sengupta, Alyson Sujkowski, Kozeta Libohova, Autumn C. Harris, Robert Wessells, Diane E. Merry, Sokol V. Todi

**Affiliations:** Department of Physiology, Wayne State University School of Medicine, Detroit, MI, USA; Department of Biochemistry and Molecular Biology, Thomas Jefferson University Sidney Kimmel Medical College, Philadelphia, PA, USA; Department of Pharmacology, Wayne State University School of Medicine, Detroit, MI, USA; Maximizing Access to Science Careers Program, Wayne State University, Detroit, MI, USA; Department of Neurology, Wayne State University School of Medicine, Detroit, MI, USA

## Abstract

Spinal and bulbar muscular atrophy (SBMA) is an X-linked disorder that affects males who inherit the *androgen receptor* (*AR*) gene with an abnormal CAG triplet repeat expansion. The resulting protein contains an elongated polyglutamine (polyQ) tract and causes motor neuron degeneration in an androgen-dependent manner. The precise molecular sequelae of SBMA are unclear. To assist with its investigation and the identification of therapeutic options, we report here a new model of SBMA in *Drosophila melanogaster*. We generated transgenic flies that express the full-length, human AR with a wild-type or pathogenic polyQ repeat. Each transgene is inserted into the same “safe harbor” site on the third chromosome of the fly as a single copy and in the same orientation. Expression of pathogenic AR, but not of its wild-type variant, in neurons or muscles leads to consistent, progressive defects in longevity and motility that are concomitant with polyQ-expanded AR protein aggregation and reduced complexity in neuromuscular junctions. Additional assays show adult fly eye abnormalities associated with the pathogenic AR species. The detrimental effects of pathogenic AR are accentuated by feeding flies the androgen, dihydrotestosterone. This new, robust SBMA model can be a valuable tool towards future investigations of this incurable disease.

## Introduction

The polyglutamine (polyQ) family of disorders comprises nine age-dependent, hereditary neurodegenerative diseases^1, 2^. Each is caused by the anomalous expansion of a CAG triplet nucleotide repeat in a disease-specific gene, with subsequent translation resulting in the corresponding protein containing an abnormally expanded polyQ tract^1^. Spinal and bulbar muscular atrophy (SBMA; also known as Kennedy’s Disease) is one member of this family of disorders. In SBMA, the CAG triplet repeat expansion resides within the *androgen receptor* (*AR*) gene on the X chromosome^3^. The other polyQ disease family members are Huntington’s disease, spinocerebellar ataxia Types 1, 2, 3, 6, 7, and 17, and dentato-rubralpallidoylusian atrophy^1^. There are currently no effective treatments for any of these diseases.

SBMA is a male-specific disease due to a requirement for circulating testosterone^4-6^. Patients experience various symptoms impacting endocrine^7, 8^, sensory^9^, metabolic^8, 10^, and motor functions^11^. Of these, the most prominent is motor impairment^12-14^. At the cellular level, polyQ-expanded AR has multiple toxic properties, including alterations in transcriptional activity^15-21^ and protein-protein interactions^22^, increased DNA-binding^15^, and impaired nuclear export^23^. PolyQ-expanded AR misfolds, forms nuclear aggregates, accumulates in inclusions, and leads to the degeneration of spinal and brainstem motor neurons^4, 24^. All of these effects culminate in slowly progressing skeletal muscle weakness and atrophy in the proximal limbs, dysarthria, dysphagia, and eventually motor dysfunction^4, 11^. Although SBMA is classically thought of as a motor neuron disease, there is strong evidence implicating the expression of AR in skeletal muscle as a contributing factor to pathology^4, 25-31^.

The AR (NR3C4) is a nuclear hormone receptor^32^ and transcription factor activated by the binding of androgens, such as testosterone and its derivative, dihydrotestosterone (DHT). When not bound to a ligand, AR resides in a cytoplasmic aporeceptor complex that includes heat shock proteins. Upon ligand binding, AR undergoes conformational changes resulting in its translocation to the nucleus. Within the nucleus, it dimerizes, binds to androgen-response elements (AREs) on DNA, and facilitates the transcription of androgen-responsive genes. Both ligand-binding and subsequent nuclear translocation are implicated in SBMA pathogenesis^6, 33-36^.

Various cell and animal models of SBMA provided valuable insights into disease mechanisms. Still, a robust *in vivo* model able to recapitulate both the physiological and biochemical aspects of the disease, while also allowing for rapid investigations of newly emerging theories, would be beneficial to the field. *Drosophila melanogaster* is a versatile model organism for studying neurodegenerative disease^37, 38^. Unlike vertebrate models, fruit flies allow for large, lifespan-long experiments to be completed in a short period of time with relatively low cost. Flies also have more complex nervous systems than yeast or cultured cell models. Additionally, because they have high conservation of human disease-associated proteins, the *Drosophila* models of progressive, age-related neuronal disorders represent a unique opportunity to study disease-induced changes in overall function.

Previous *Drosophila* models of SBMA^33, 34, 39-42^ were generated using the modular GAL4/UAS system^43^. These models expressed either a truncated^41^ or full-length form of human polyQ-expanded AR protein^33, 42^ in a tissue-specific manner. Tissues targeted for polyQ-expanded AR included the eye^33, 34, 41^, salivary gland^34^, central and peripheral nervous systems^41^, and motor neurons^34, 41^. These models demonstrated ligand-dependent degeneration^34^, motility defects^34, 41^, alterations in larval neuromuscular junction (NMJ)^34^, and shortened lifespan^34, 41^. To align with existing mammalian models and human disease, we developed new *Drosophila* models of SBMA that express full-length, human AR in both the WT and the pathogenically expanded-CAG repeat form. Here, we provide the initial characterizations of the new SBMA lines with emphasis on phenotypes particularly relevant to human disease; longevity, motility, and NMJ pathology.

## Results

### Generation of WT and polyQ-expanded AR Drosophila lines

To generate the new SBMA lines, cDNA sequences encoding the full-length human AR with either wild-type (Q20) or expanded (Q112) repeats (Figure 1A) were inserted into a safe harbor locus using pWalium.10.moe as a targeting vector (Figure 1B). To minimize the possibility of additional toxicity from mRNA or unconventional translation processes, as well as triplet repeat-associated instability^44-48^, our models contain a mixed CAA/CAG repeat sequence. The presence of AR protein was visualized via Western blotting (WB) (Figure 1C). The main wild-type (WT) and polyQ-expanded (SBMA) AR bands (black arrows) migrate above the 100 kDa marker, with the expanded variant running higher due to the longer polyQ (Figure 1C). Also of note is the presence of SDS-resistant, polyQ-expanded species in the stacking (solid-red bracket) and resolving (dotted-red bracket) portions of the gel, as well as lower molecular weight bands that are likely proteolytic products of AR (gray bracket).

**Figure 1:**
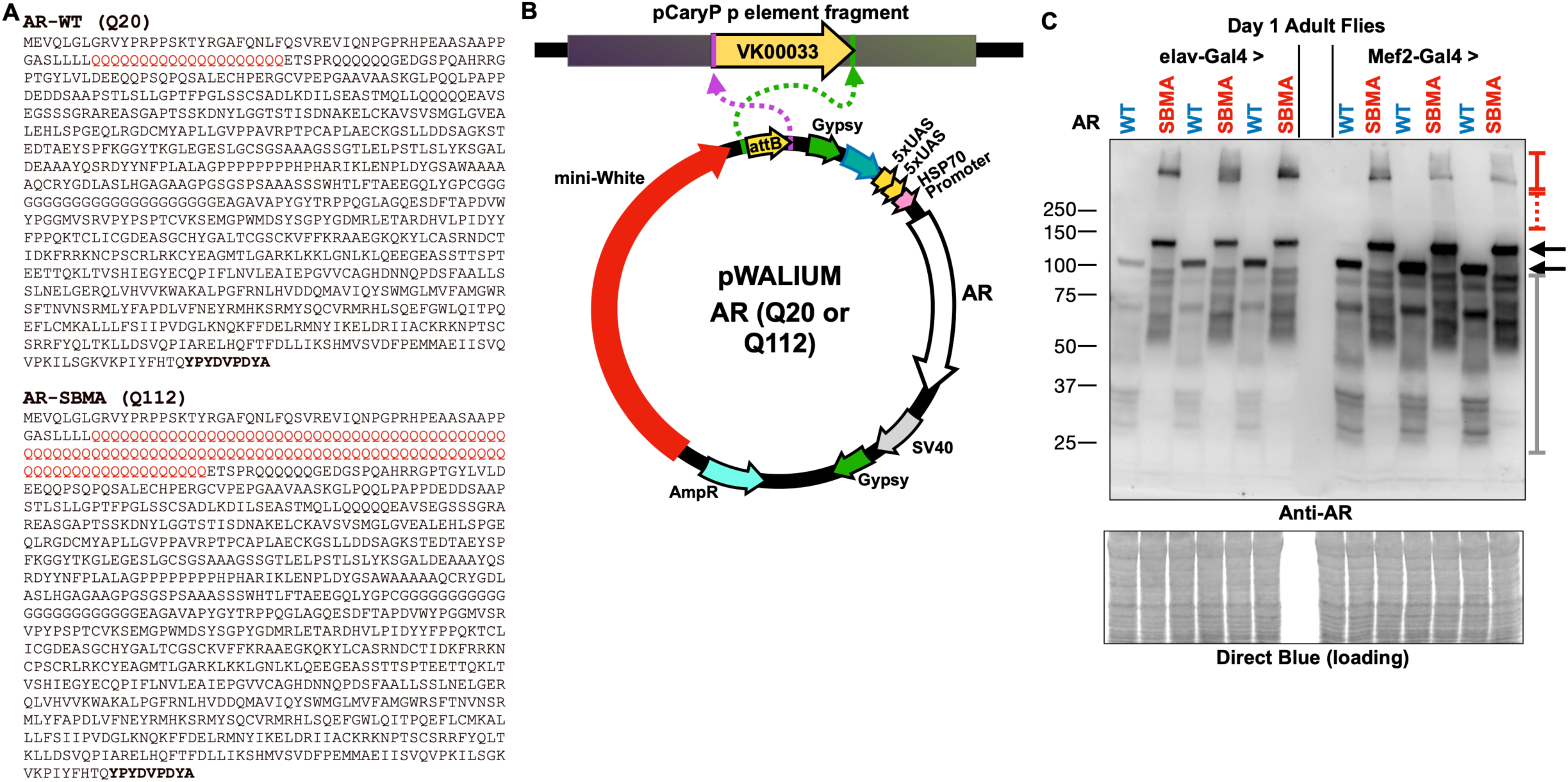
New transgenic fly lines for SBMA. A) Amino acid sequence of the human AR protein used. The polyQ is encoded by a CAGCAA doublet repeat to circumvent the possibility of mRNA- and non-AUG translation-dependent toxicity. Bolded: HA epitope tag. B) Insertion strategy into site VK00033 on chromosome 3 of the fly. C) Western blotting from adult flies expressing the noted transgenes. Whole fly lysates. elav-Gal4: pan-neuronal expression. Mef2-Gal4: muscle cell expression. Black arrows: main AR bands; solid red bracketed line: SDS-resistant, polyQ-expanded AR trapped in the stacking gel; dotted, red bracketed line, polyQ-expanded AR trapped in resolving gel; gray bracketed line: likely proteolytic products of AR. Each lane is from an independent repeat.

### Expression of polyQ-expanded AR in neurons or muscle reduces lifespan and impairs mobility in adult Drosophila

To characterize the phenotypes of these lines, we examined if expression of WT or SBMA AR causes deficits in the absence of ligand. Because AR toxicity affects both motor neurons and skeletal muscle^4, 11^, we utilized the Gal4/UAS system to drive AR in the nervous system or muscles in flies. This modular system utilizes the exogenous yeast transcriptional activator Gal4 and its binding sequence, UAS (<u>U</u>pstream <u>A</u>ctivating <u>S</u>equence) to manipulate expression of target transgenes in specific tissues within the fly throughout development and adulthood^43, 49^. By mating one fly containing Gal4 under the control of a tissue-specific promotor (driver) and one fly containing the UAS (responder) sequence upstream of the target gene, transcription can be targeted to the tissue of interest. Myocyte enhancer factor-2 (Mef2) is required for *Drosophila* muscle differentiation and development and is expressed in both cardiac and skeletal muscle^50-52^. Expressing Gal4 under the control of the Mef2 promoter (Mef2-Gal4) allows for muscle-specific expression of target transgenes. Embryonic lethal abnormal vision (elav) is expressed pan-neuronally and is required for developing and maintaining the *Drosophila* nervous system^53, 54^.

We found that in the absence of ligand, SBMA AR dramatically reduced longevity with both neuronal and muscle tissue expression (Figure 2A). Expressing WT AR in neuronal cells also resulted in modest reductions to longevity (Figure 2A). However, the reduction in longevity with WT AR appears to be inconsistent, as it is absent in other repetitions (please refer to the “EtOH” group in Figure 4A). In muscles, WT AR expression slightly, but significantly, increased longevity (Figure 2A; the same pattern was independently confirmed in Figure 5A); the cause of this modest increase is unclear and will be a focus of future investigations. In assessments of climbing ability, SBMA flies became impaired significantly more quickly than control flies (Figure 2B).

**Figure 2:**
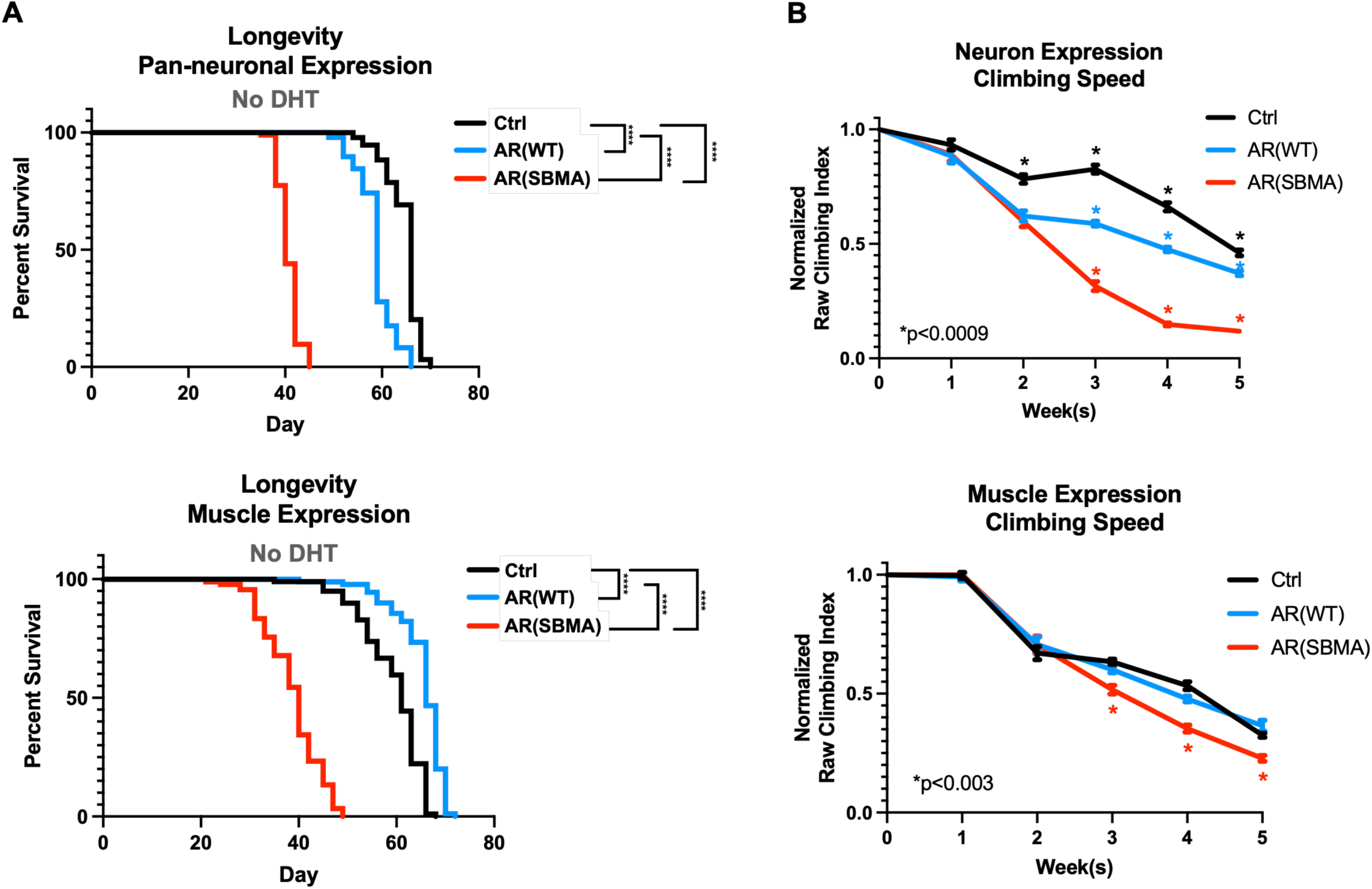
Longevity and motility outcomes from expression of WT or polyQ-expanded AR in neuronal or muscle cells. A) Longevity assays of female flies expressing the noted transgenes. No DHT was introduced at any point of their lives. N≧100 flies per group. ****: p<0.0001, Log-rank tests. B) Normalized motility assays of female flies expressing the noted transgenes in neurons (N) or muscles (M). N≧100 flies per group. Statistical analyses used: 2-way ANOVA with post hoc correction, mean ± SEM. *: p<0.0009 in neurons, p<0.003 in muscle. For (A) and (B), Ctrl denotes the background line used to generate the transgenic lines, in trans with the respective driver.

**Figure 3:**
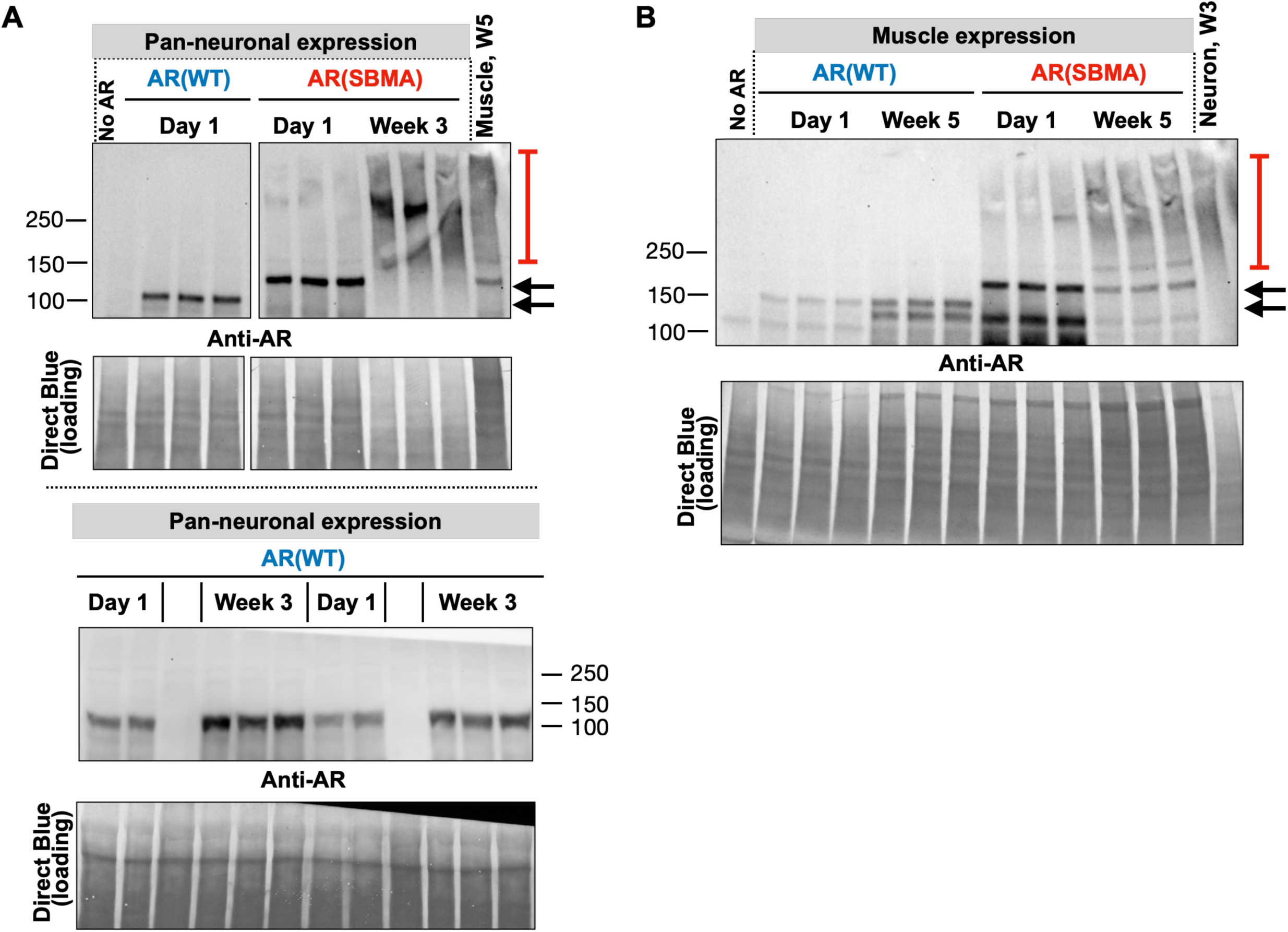
Expression of AR protein in neurons and muscles. Western blots from whole flies expressing the noted transgenes in neurons (A) or muscles (B), for the amounts of time indicated. Images in top part of panel (A) are from the same blot and exposure, cropped and rearranged to ease visualization. Blots in bottom part of panel (A) are from independent lysates. In both panels, each lane is an independent repeat, i.e., for (A) N=3 for top panel, except for the first lane without AR and the last lane, with muscle expression, where N=1; and N=4 for day 1 and N=6 for week 3 on the bottom panel; for B: N=3 for all lanes, except first and last lanes, where N=1.

**Figure 4:**
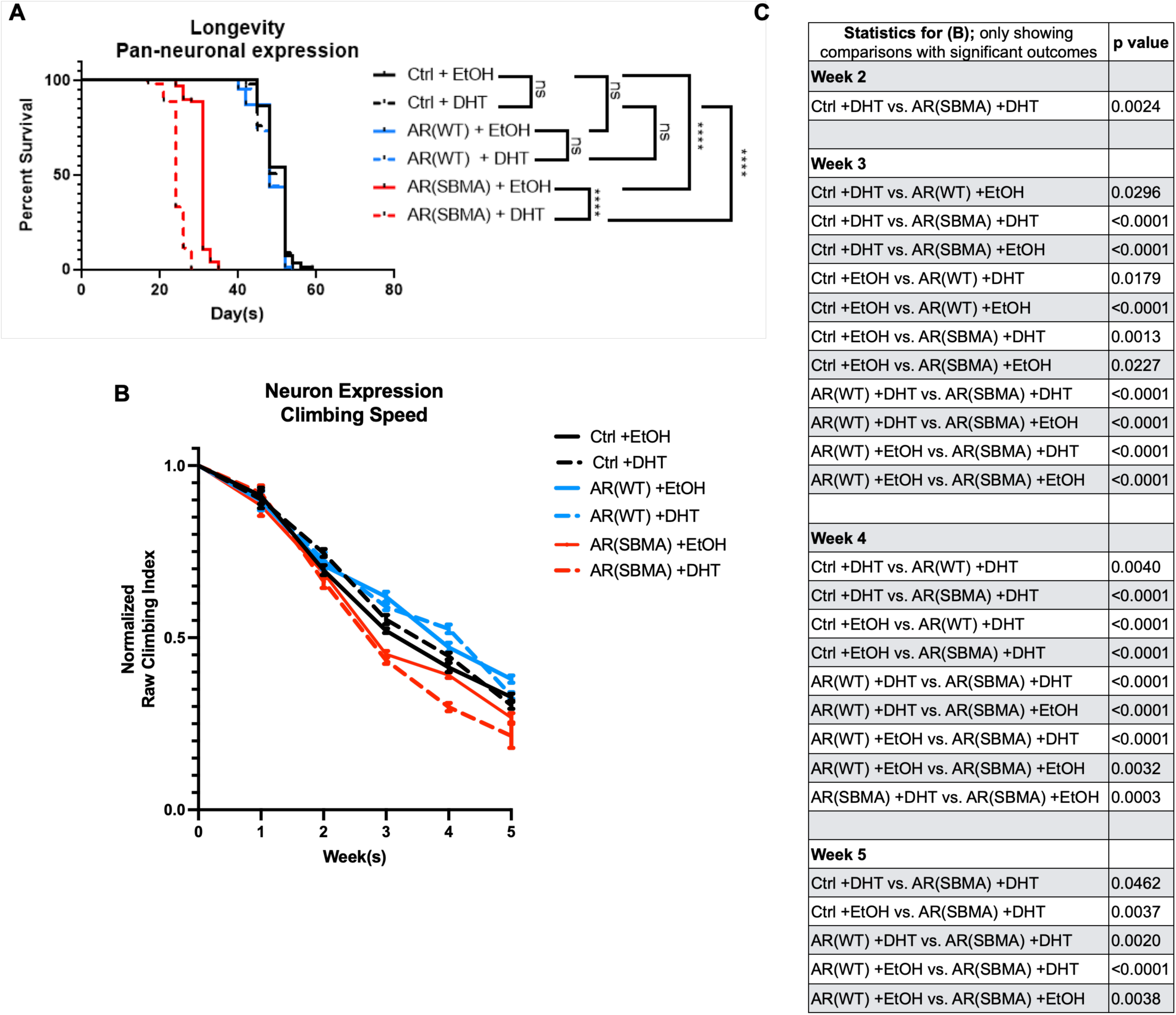
Effect of DHT on SBMA model motility and longevity when expressed in neurons. (A) Longevity and (B) Normalized motility outcomes from expression of normal or polyQ-expanded AR in neuronal cells, without or with DHT. N≧100 flies per group for (A) and (B). Statistical analyses used: Log-rank tests, ****: p<0.0001 (longevity), 2-way ANOVA with post hoc correction, mean ± SEM, Statistics summarized in (C) (motility).

**Figure 5:**
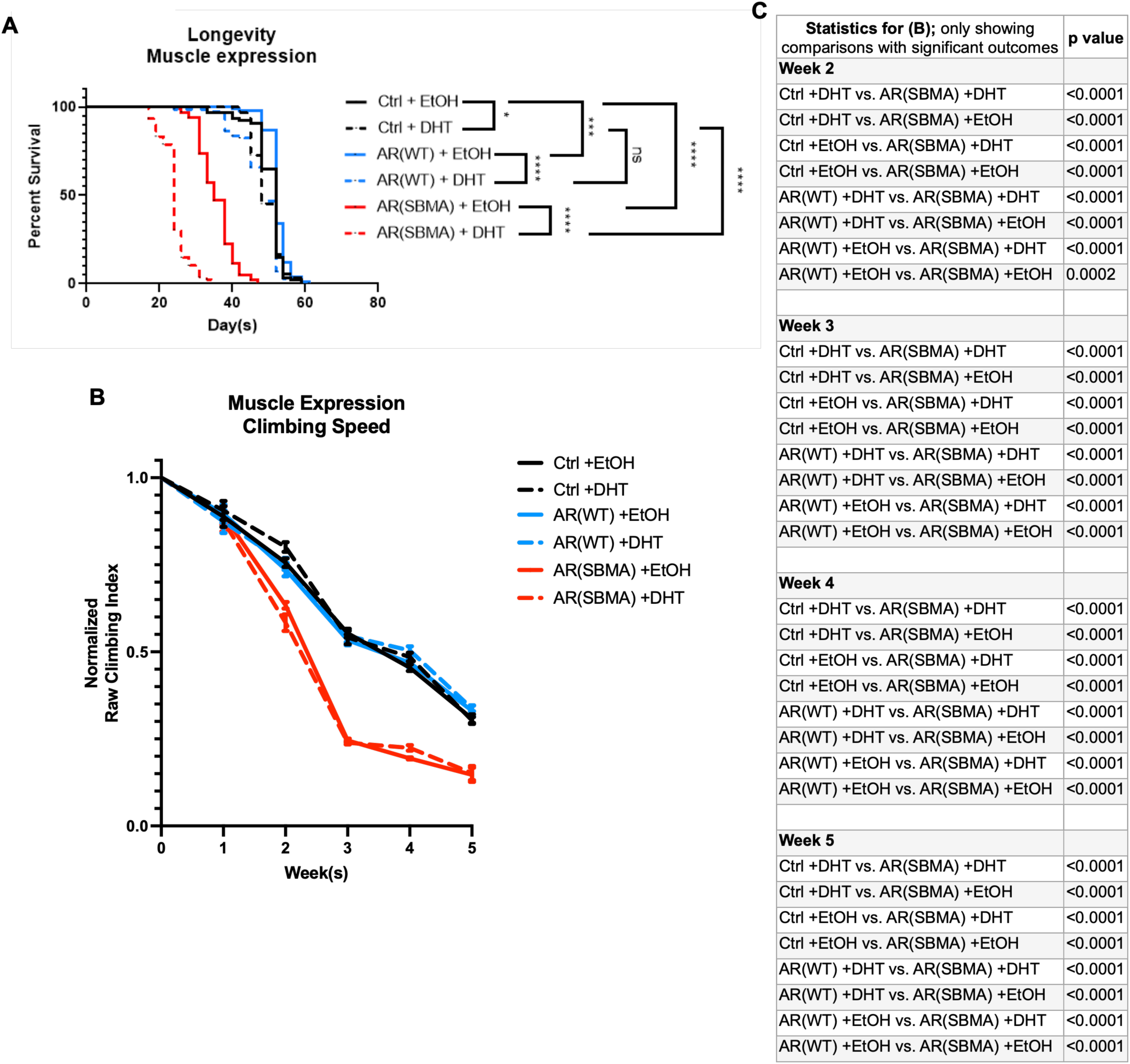
Effect of DHT on SBMA model motility and longevity when expressed in muscles. (A) Longevity and (B) Normalized motility outcomes from expression of normal or polyQ-expanded AR in neuronal cells, without or with DHT. N≧100 flies per group for (A) and (B). Statistical analyses used: Log-rank tests, *: p<0.05, ***: p<0.001, ****: p<0.0001 (longevity), 2-way ANOVA with post hoc correction, mean ± SEM, Statistics summarized in (C) (motility).

Interestingly, although the effect was milder than with SBMA AR, pan-neuronal expression of WT AR also reduced climbing ability compared to controls lacking AR. This decline-of-motility phenotype was absent in flies with muscle expression of AR (Figure 2B). At first, these data showing negative phenotypes in the absence of DHT were surprising since polyQ-expanded AR toxicity requires ligand-activation. However, our findings are in agreement with more recent reports of some androgen-independent pathology in disease conditions^55^.

The experiments in figures 1 and 2 were conducted using female flies. Male flies also showed marked reductions in longevity with neuronal and muscle expression of polyQ-expanded AR, while robust motor deficits were seen only with neuronal expression (Supplemental Figure 1). For additional characterizations, we elected to move forward focusing only on female flies. Even though SBMA is male-specific in humans, AR is exogenous to both male and female flies; therefore, the choice to examine female flies in this context rather than males does not lessen its relevance to the human disease. Moreover, the use of male mice is necessary in mammalian models of this disease, because they produce androgens; flies do not produce testosterone or DHT and since SBMA is hormone-dependent, the flies in the study do not need to be male. Additional rationale for this choice is as follows: using one sex reduces the overall amount of DHT required for the study, thus minimizing the use of a controlled substance; since both sexes showed strong phenotypes with expression of polyQ-expanded AR, utilizing one sex still provides pertinent information that can be extrapolated to both; lastly, for the purposes of histological and biochemical preparations, the larger size of female flies compared to males yields modestly higher amounts of material for downstream applications and eases anatomical dissections and observations.

To visualize the relationship between physiological outputs and AR protein levels and aggregation, we next conducted WBs. We selected two time points for each expression pattern: day one and week three for neurons, and day one and week five for muscle. We selected the three-week time point for neuronal expression as it coincides with sustained, significant reduction in mobility for this tissue type; we chose the five-week mark for muscle expression for the same reason. As shown in figure 3, expression of AR was consistent during this time frame, in each tissue. With the disease-causing version, we observed the accumulation of SDS-resistant smears over time, concomitant with a reduction in signal of the primary AR band. This effect was especially pronounced in neuronal tissue.

From these results (figures 1-3), we conclude that expression of polyQ-expanded AR is toxic when expressed in fly neuronal and muscle tissue and that its toxicity coincides with increased aggregation of the insulting protein.

### DHT reduces longevity and exacerbates motor impairment in SBMA model flies

Because SBMA is an androgen-dependent disorder^4, 5^, we next evaluated the role of DHT in the longevity and motility of WT and SBMA AR flies. With pan-neuronal expression, DHT supplementation had no significant effect on the longevity of either the background control or WT AR flies (Figure 4A). In agreement with non-DHT studies, polyQ-expanded AR expression in neurons resulted in severe reduction to lifespan, further enhanced by DHT supplementation (Figure 4A). These toxic effects carried over to mobility, where SBMA lines rapidly declined over time, a phenotype that was again exacerbated by DHT (Figure 4B).

Unlike with neuronal expression, DHT supplementation with muscle expression of AR negatively impacted the lifespan of both WT AR and SBMA AR flies (Figure 5A), although this phenotype was more severe with SBMA AR. The reduction in longevity observed in flies not expressing AR in the presence of DHT was not a consistent outcome, as we did not always observe a significant reduction in lifespan in these flies (supplemental figure 2). Muscle expression of polyQ-expanded AR consistently reduced lifespan both with and without the addition of DHT. However, this effect was worsened by DHT supplementation (Figure 5A). In assessments of mobility, as observed in Fig. 2B, SBMA lines exhibited significantly reduced climbing ability starting in week two and continuing through week five (Figure 5B). Interestingly, DHT did not impact the rate of decline in climbing ability for any of the lines tested (Figure 5B).

We next examined the effect of DHT on WT and SBMA AR using WBs. We focused again on week three for neurons and week five for muscles, similar to the experiments conducted in the absence of DHT (Figure 3). As reported in other models^56, 57^, DHT led to increased levels of AR protein in both WT and SBMA models, in both neurons and muscles (Figure 6). With polyQ-expanded AR, we again observed the presence of SDS-resistant species regardless of the treatment with vehicle control or DHT, but the types of smears seen were different, indicating differences in the types of aggregates formed in the presence of ligand. We also noticed trends that did not reach statistical significance with the levels of SDS-resistant species in the absence *versus* presence of DHT: higher levels with SBMA AR in neurons and muscles (figure 6), and reduced (neurons; figure 6A) or similar levels (muscle; figure 6B) with WT AR.

**Figure 6:**
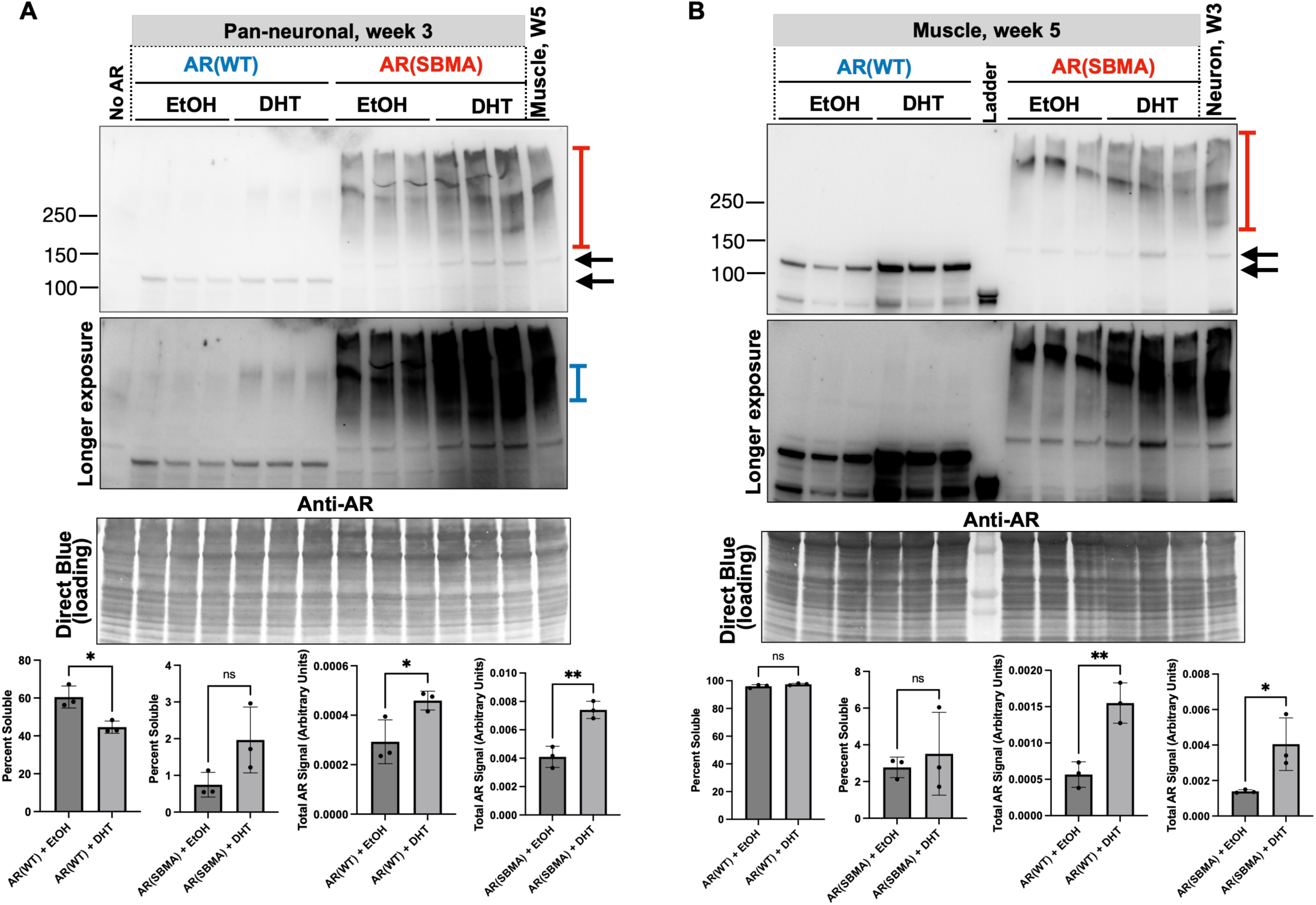
AR protein in the absence and presence of DHT. Western blots from whole flies expressing the noted AR proteins. Black arrows: main AR bands; red bracketed lines: SDS-resistant, polyQ-expanded AR; blue bracketed arrow, SDS-resistant, WT AR. Quantifications are from the images on top, where each lane represents an independent repeat. *: p<0.05; **: p<0.01 based on Student’s t-tests. Each lane is an independent repeat. Shown in graphs are means ± SD.

Altogether, these data indicate that DHT worsens phenotypes caused by polyQ-expanded AR when expressed in neurons or in muscles, in agreement with reports from other models of SBMA^33, 34, 36, 58^.

### The effect of SBMA AR on neuromuscular junction complexity

In mouse and previous fly models of SBMA, the structure and function of the neuromuscular junction (NMJ) is pathologically altered^16, 34, 59-62^. Here, we examined NMJ complexity in the dorsal longitudinal flight muscles to assess the effect of polyQ-expanded AR expression in our fly models (Figure 7, Supplemental Figure 3). On day one, no significant differences in NMJ complexity were observed with pan-neuronal or muscle expression of either AR transgene (Figure 7). At three weeks of age, pan-neuronal expression of SBMA AR significantly reduced NMJ complexity, with or without DHT supplementation; DHT led to a trend of further reduction in complexity, but this did not reach statistical significance (Figure 8A,B). Interestingly, the NMJ complexity of WT AR flies also trended lower with the addition of DHT (Figure 8A,B); however, this also did not reach statistical significance. We did not observe significant differences in NMJ complexity at five weeks of age in flies with muscle-specific expression of WT or polyQ-expanded AR (Figure 8C,D).

**Figure 7:**
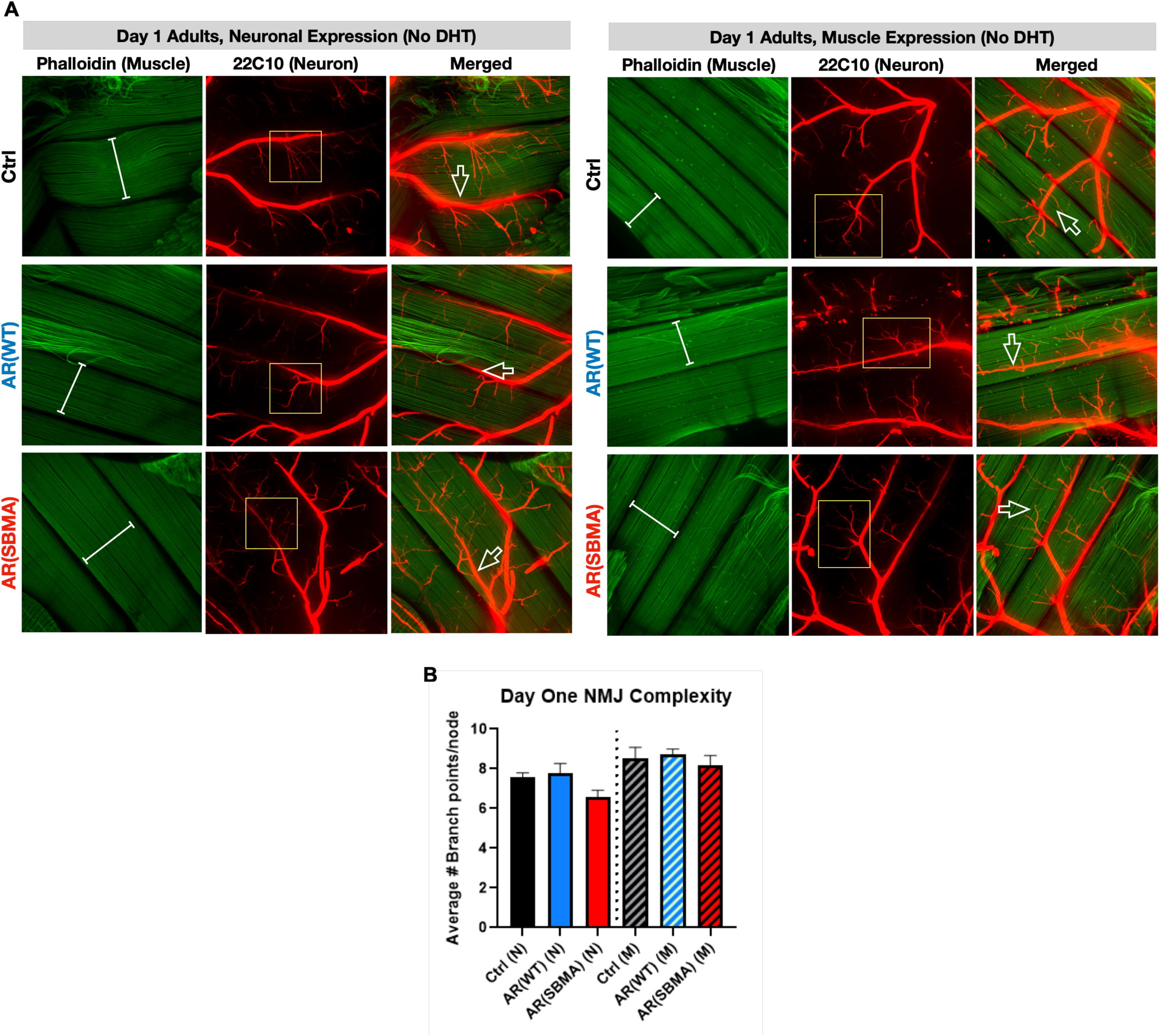
Effects of AR on neuronal-muscle structural relation in young flies. A) Representative images from adult female flies whose flight muscles were dissected and stained as outlined in methods. White bracketed arrows: muscle fibers; open arrows: motoneuron branches that are adjacent to muscle fibers; yellow boxes: neuronal branching. B) Quantification of axonal branching complexity separated by expressing tissue. Statistics: not significant by one-way ANOVA. N≧12 muscles per condition.

**Figure 8:**
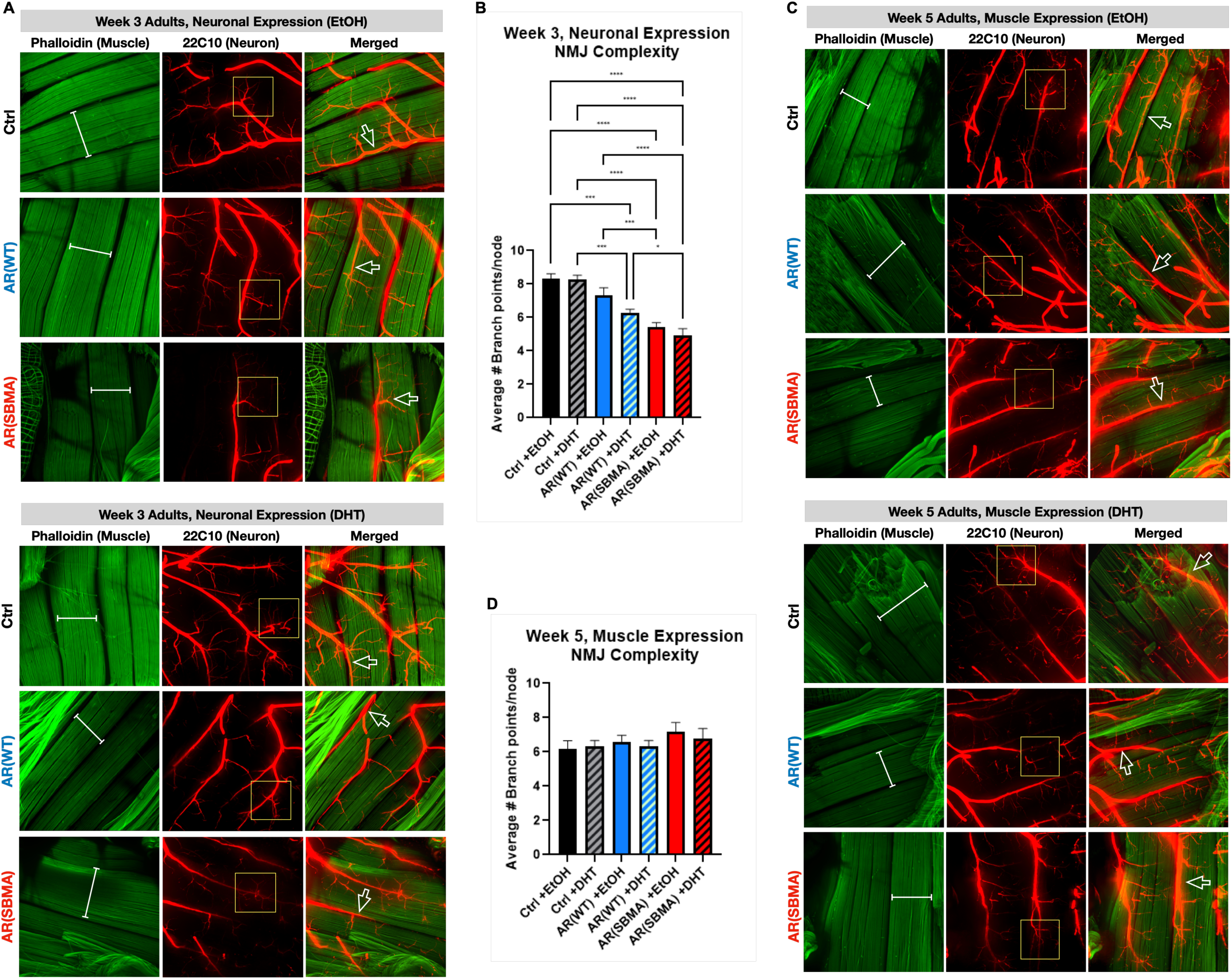
Effects of AR expression on neuronal-muscle structural relation in aged adult flies with or without ligand. A) Representative images from adult female flies whose flight muscles were dissected and stained as noted. Pan-neuronal expression of WT or SBMA AR with DHT or EtOH. White bracketed arrows: muscle fibers; open arrows: motor neuron branches that are adjacent to muscle fibers; yellow boxes: neuronal branching. B) Quantification of axonal branching with pan-neuronal expression of AR. N≧12 muscles per condition. Statistics: two-way ANOVA with post hoc correction, mean ± SD. *: p<0.05, ***: p<0.001, ****: p<0.0001. C) Representative images from adult female flies whose flight muscles were dissected and stained as noted. Muscle expression of WT or SBMA AR with DHT or EtOH. White bracketed arrows: muscle fibers; open arrows: motoneuron branches that are adjacent to muscle fibers; yellow boxes: neuronal branching. D) Quantification of axonal branching with pan-neuronal expression of AR. Statistics: not significant by two-way ANOVA. N≧12 muscles per condition.

### ARQ112 causes deterioration in Drosophila eye

Eye-specific expression of mutant genes in *Drosophila* is a well-established tool in the study of neurodegenerative diseases – it is useful for conducting large genetic screens due to the ease of phenotype evaluation. Moreover, mutations that would be lethal in other cell types (e.g., neurons, or muscles) can be assessed in fly eyes without impacting viability. Previous work established a useful scoring system (illustrated in figure 9A) for tracking eye deterioration over time^63, 64^. We utilized this scoring system by crossing each of our lines to the eye-restricted driver, GMR-Gal4.

**Figure 9:**
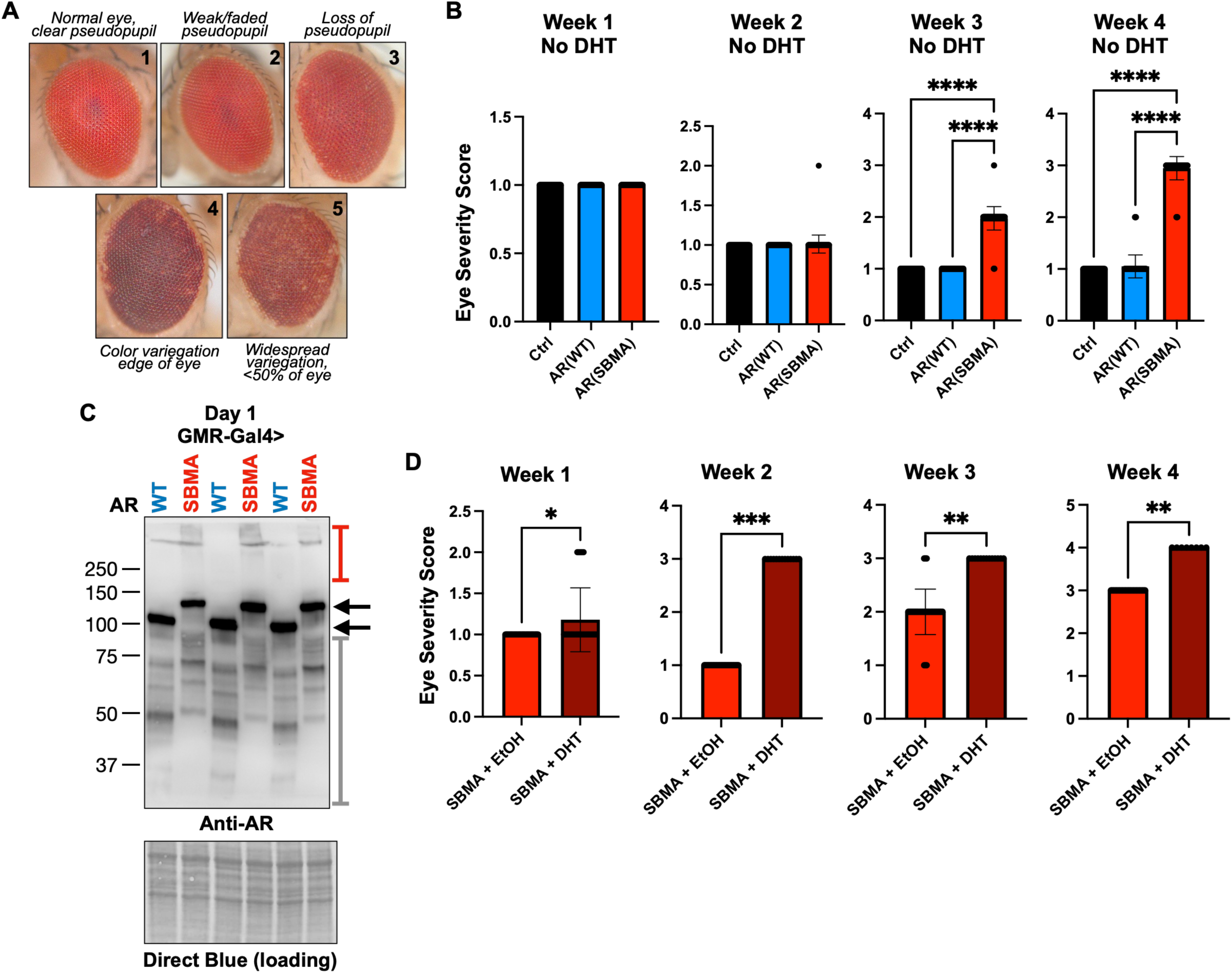
Effects of AR expression in fly eyes. A) Scoring system. B, D) Impact of AR expression in fly eyes for the noted days, in the presence or absence of DHT. WT: wild-type AR. SBMA: polyQ-expanded AR. Means +/-SD. Statistics: Kruskal-Wallis tests, where *: p<0.05, **: p<0.01, ***: p<0.001, ****: p<0.0001. N≧42 per group per time point in (B) and N≧8 (D). For (D), we kept the “N” small to highlight the significance of the phenotype. C) Western blots from dissected fly heads expressing the noted forms of AR in eyes in the absence of ligand. Each lane represents an independent repeat. Black arrows: main AR bands; red bracketed line: SDS-resistant AR; gray bracketed line: likely proteolytic fragments of AR protein.

In the absence of DHT, SBMA AR expression worsened eye phenotypes beginning in week three, while WT AR and non-AR expressing controls remained unaffected throughout the timeline of investigations (Figure 9B). Examination via WBs showed a pattern of migration for WT and SBMA AR similar to what we observed in other tissues, with SBMA AR migrating as a main band as well as SDS-resistant species (Figure 9C). The SBMA AR-dependent phenotype in fly eyes worsened with the addition of DHT, being exacerbated as early as the second week and continuing through week four (Figure 9D). Thus, as with other tissues that we examined, SBMA AR also causes pathology in the fly eye.

## Discussion

We sought to generate robustly phenotypic *Drosophila* models of SBMA that can help the field with expedited investigations of the biology of this disease and the identification of therapeutic options. The experiments summarized here demonstrate the value of the new SBMA models. In agreement with data from mammalian models of this disease, the new *Drosophila* SBMA lines have reduced lifespan, motility, and NMJ complexity, coinciding with aggregation of polyQ-expanded AR protein. These phenotypes are generally worsened by the addition of DHT. In line with our data, DHT-dependent lethality and motor impairment have also been reported in previous adult fly and larval models expressing human full-length polyQ-expanded AR^16, 34, 65^. When expressed in neurons, SBMA AR consistently and rapidly reduced each of the physiological outputs that we measured and led to the deterioration of NMJ complexity. DHT-dependent reduction in NMJ branching complexity was previously reported in fly larvae expressing human polyQ-expanded AR in a motor neuron-specific manner^34^. In our model, we did not observe this difference at the time points we examined. We chose histology timepoints based on the strong phenotypic deficits that we observed, rationalizing that they would concur with NMJ alterations; however, it is possible that there are significant NMJ changes between the treatments that were not captured by this specific choice of timeline. When expressed in muscles, SBMA AR again led to general reductions in physiological outputs; however, these did not coincide with structural deteriorations in the NMJ. Alongside reports from studies in mammals, these data support important roles for both neuronal and muscle tissue in future investigations of SBMA biology of disease.

A key feature of SBMA pathogenesis is the ligand-enhanced translocation of polyQ-expanded AR to the nucleus. Contrary to other *Drosophila* models^33, 34, 42^, expressing human polyQ-expanded AR in our fly models resulted in marked phenotypes in the absence of ligand. In our ongoing work, we noticed that both WT and SBMA AR indeed localize to the nucleus in fly tissues in the absence of DHT; presence in the nucleus is enhanced by the supplementation of DHT for SBMA AR, whereas WT AR shows a trend of increased nuclear fractionation, but does not reach statistical significance (Supplemental Figure 4). This is distinct from earlier studies in PC12 cells in which polyQ-expanded AR aggregation and toxicity is completely ligand dependent^36, 58^.

Moreover, forced localization of the expanded AR to the nucleus did not result in its aggregation in the absence of ligand^36^. Only when the AR was activated (conformationally altered) by ligand binding, did its toxic effects ensue. However, our observations of ligand-independent phenotypes with SBMA AR are congruent with recent studies in transgenic mice, which suggested that early events in SBMA can be androgen-independent^61^. Moreover, it is known that AR can be imported into the nucleus in the absence of androgens^66^, (although even in this location, it is largely inactive in the absence of ligand.).

Ligand-free, AR-induced toxicity at the level of the mitochondria was recently reported in patient-derived cells^55^; ligand-binding did exacerbate toxicity^55^. It is of interest that mixed or alternating CAG/CAA repeats were used in the development of these cell lines^55^, similar to the design of our models. Pure *versus* mixed polyQ-encoding repeats lead to different types of mRNA structures: pure CAG repeats result in relatively stable hairpins, while mixed CAG/CAA repeats result in unstable hairpins^45^. One possibility is that the different mRNA structures formed by polyQ-encoding repeats lead to mRNA-based toxicity at the motor unit. mRNA toxicity has been implicated in various triplet repeat diseases, although that toxicity usually has been associated with pure CAG repeats, rather than mixed ones^67^. Whether such unstable, or reduced, hairpin formation enhances mixed CAG/CAA repeat translation efficiency, leading to its enhanced nuclear localization and aggregation, or an RNA-mediated mechanism of toxicity, is the focus of ongoing investigations.

An additional point of interest is the negative impact of WT AR overexpression in previous *Drosophila* models of SBMA^17, 34^. These findings are consistent with studies in mice, where substantial overexpression of WT AR in muscle resulted in severe phenotypes^26^. Because there appear to be expression-level dependent influences on phenotype severity, it may be that higher expression of WT AR protein amplifies native AR functions and contributes to toxicity^34^. In our model, we saw minor or no phenotypes with WT AR expression in the tissues we examined. In some cases, the phenotypes seen with WT AR were actually improved outcomes, compared with controls (Figure 2A; longevity with muscle expression). However, these differences were inconsistent and varied with age, leading us to conclude that overexpression of WT AR in our fly models is generally non-toxic.

To conclude, we introduce a new model of SBMA in *Drosophila melanogaster*. This model recapitulates key phenotypic manifestations of the disease in neuronal and muscle tissues and can be used for hypothesis-based examinations or screening efforts to help further our understanding of this disorder and the discovery of therapeutic interventions for it.

## Materials and Methods

### Fly stocks and maintenance

All flies were aged at 25°C under controlled 50% humidity and 12-hour light/dark cycle. Virgin females and males used for crosses and experiments were collected under light CO2 within 2 hours of eclosion over a 72-hour period. All flies used in experiments were age matched and heterozygous for driver and responder. Stocks used for experiments were obtained as follows: GMR-Gal4 from BDSC (Bloomington Drosophila Stock Center) (Stock #8605), elav-Gal4 was generously gifted by Dr. Daniel Eberl, University of Iowa, and Mef2-Gal4 from Dr. Rolf Bodmer, Sanford-Burnham Medical Research Institute, California.

### New fly line generation

ARQ20 and ARQ112 cDNA sequences were based on the NCBI human AR reference sequence NM_000044.2. Mutagenesis was carried out by Genscript (genscript.com) and codon-optimized for expression in *Drosophila*. The transgenes were subcloned into pWalium10-moe and fly lines were generated using pHiC31-dependent integration into VK00033 on chromosome 3. All resulting transformants were migrated into the w^1118^ background.

### DHT Treatment

Flies used for ligand experiments were collected and then divided at random into cohorts of DHT- or EtOH (vehicle)-treated. After being separated, DHT flies were transferred onto vials containing 80uL of 20mM DHT dissolved in 100% EtOH, while EtOH controls were transferred into vials containing 80uL of 100% EtOH. DHT or EtOH was pipetted directly on top of the food and allowed to absorb for 1 hr prior to use. Fresh vials were provided three times each week for the duration of the study. The first day of DHT treatment was considered “Day One” in all DHT experiments. DHT was procured from Cayman Chemicals (cat. # 15874).

### Rapid negative geotaxis speed assay

Climbing speed was assessed in Rapid Integrative Negative Geotaxis (RING) assays using groups of 100 flies per cohort. Five vials containing 20 flies each were set-up in a RING apparatus and negative geotaxis stimulus was initiated by tapping the apparatus down on the countertop twice firmly and rapidly. The height of the flies in the vials were captured by photo image 2 seconds after the start of climbing. Flies were longitudinally tested twice per week for 5 weeks to assess climbing ability at each aged time point. Longitudinal climbing speed is normalized to day one climbing data within each genotype and treatment. For descriptions in further detail refer to Damschroder *et al.* (2018)^68^.

### Longevity

Differences in survival were measured using cohorts of 100 sex- and age-matched flies per condition. Vials were inspected and changed every other day to count and remove any dead flies. Flies that escaped the vial or died for reasons other than natural aging were excluded from analysis.

### NMJ Isolation and Staining

NMJ dissections and staining were modified from Sidisky *et al*. (2020)^69^. Briefly, whole flies were anesthetized in light CO2 then submerged for 30 seconds in 70% ethanol to remove wax coating from cuticle before being transferred to a gel bottomed dissecting dish. The wings, legs, head, and abdomen were removed, and thoraxes were transferred to 4% paraformaldehyde for 60 minutes. Fixed thoraces were submerged in liquid nitrogen for 10 seconds, bisected with a sharp razor blade under a dissecting scope, and transferred to ice cold PBS. Hemithoraces were then blocked for two hours before staining overnight with anti-22c10 primary antibody (mouse a-22c10, 1:100, DSHB). The following day, tissues were washed, and stained with conjugated primary and secondary antibodies (a-mouse AF594, 1:200, ThermoFisher and phalloidin AF488, 1:500, ThermoFisher), and imaged using a confocal microscope. Each of the data points used for quantifications of NMJ complexity were provided as the average number of branch points of 4 individual nodes within a single muscle prep for each fly analyzed (supplemental figure 3). Images were blinded prior to analysis.

### Western Blots

Western blotting was performed with 5 whole adult flies (neuronal or muscle expression) or 10 dissected fly heads (eye expression) per sample. Each sample was physically homogenized in 95°C lysis buffer (50 mM Tris pH 6.8, 2% SDS, 10% glycerol, 100 mM dithiothreitol (DTT)), sonicated, boiled for 10 minutes, and then centrifuged for 10 minutes at room temperature at 13,300 rpm. Homogenized samples were electrophoresed through 4-20% Tris/Glycine gels (Bio-Rad). ChemiDoc (Bio-Rad) was used to image Western blots, which were then quantified with ImageLab (Bio-Rad).Direct Blue staining of the total protein signal was used as loading control. Direct blue staining was performed as follows: PVDF membranes were incubated for 10 minutes in 0.008% Direct Blue 71 (Sigma-Aldrich) in 40% ethanol and 10% acetic acid, rinsed with 40% ethanol/10% acetic acid, air dried, and then imaged.

### Eye Scoring

Eye scores were represented in a scoring system, with higher numbers indicating worsening phenotypes characterized as follows:

1) normal (wild-type-looking) eye with a clear pseudopupil; 2) faded pseudopupil that has begun to lose its clarity; 3) no visible pseudopupil; 4) color variegation/depigmentation at the edge of the eye in addition to pseudopupil loss; 5) depigmentation throughout the eye in addition to pseudopupil loss

**Statistics:** Prism 9 (GraphPad) was used for graphics and statistical analyses. Statistical analyses used are noted in the figure legends.

## Author contributions

KR: conceptualization, data curation, software, formal analysis, validation, investigation, visualization, methodology, and writing and editing.

MS: data curation, software, formal analysis, validation, investigation, visualization, methodology, and writing and editing.

ALS: validation, investigation, visualization, methodology. KL: validation, investigation, visualization, methodology. ALH: validation, investigation, visualization, methodology.

RJW: conceptualization, data curation, funding acquisition, methodology, and writing and editing.

DEM: conceptualization, data curation, funding acquisition, methodology, and writing and editing.

SVT: conceptualization, data curation, funding acquisition, software, formal analysis, validation, investigation, visualization, methodology, and writing and editing.

