## Supplemental Figures and legends for "A phenotypically robust model of Spinal and Bulbar Muscular Atrophy in *Drosophila*"

### Supplementary information

#### Supplemental figure 1: Longevity and motility outcomes from expression of WT or SBMA AR in male flies.

Male *Drosophila* show robust lifespan and climbing speed phenotypes when expressing SBMA AR, similar to females. N≧100 flies per group for longevity and N≧100 flies per group for motility. Statistical analyses used: Log-rank tests, ****: p<0.0001 (longevity), one-way ANOVA with post hoc correction, mean ± SD, *: p<0.05, **: p<0.01, ****: p<0.0001 (climbing speed). Also shown are normalized longitudinal climbing speed. Statistical analyses used: 2-way ANOVA with post hoc correction, mean ± SEM. N≧100 flies per group. *: p<0.0001 in neurons and in muscle.

#### Supplemental figure 2: Differential effects of DHT supplementation on flies without AR transgene.

DHT supplementation effects on longevity in control flies not expressing AR transgenes were repetition dependent. Sometimes DHT had no effect on lifespan as shown here and other times DHT supplementation had a mild effect (Figure 5A). N≧100 flies per group. Not significant by Log-rank tests.

#### Supplemental figure 3: NMJ complexity quantification method.

NMJ complexity was calculated as an average of branching points per four nodes within each muscle dissection. Branch points (red arrows) were counted using the point tool in ImageJ. Right panel: the color of the dots is an automatically added feature of ImageJ and is not factored into the computations.

#### Supplemental figure 4: Subcellular fractionation of AR in adult neurons.

Western blots from whole adult flies expressing WT (A) or SBMA (B) AR in neurons for three weeks, processed to separate cytoplasmic and nuclear fractions. Quantifications are from images shown, normalized to their respective loading control (Direct Blue staining for total protein signal). Both the AR main band smears, where present, were quantified. Statistics: Welch's t-tests, where *: p<0.05, **: p<0.01, ***: p<0.001, ****:

$p<0.0001$. Shown in graphs are means ± SD. Letters at the bottom of images indicate independent repeats for each fraction. For statistics, nuclear "A" was compared to cytoplasmic "A", etc., within each genotype. Black arrow: main AR band. Red brackets: SDS-resistant AR. Fractionation was performed using the ReadyPrep Protein Extraction Kit (1632089, Bio-Rad) using seven whole flies per group that were lysed in cytoplasmic extraction buffer (Bio-Rad) and processed as delineated by the supplier's protocols.

### Supplemental figure 1

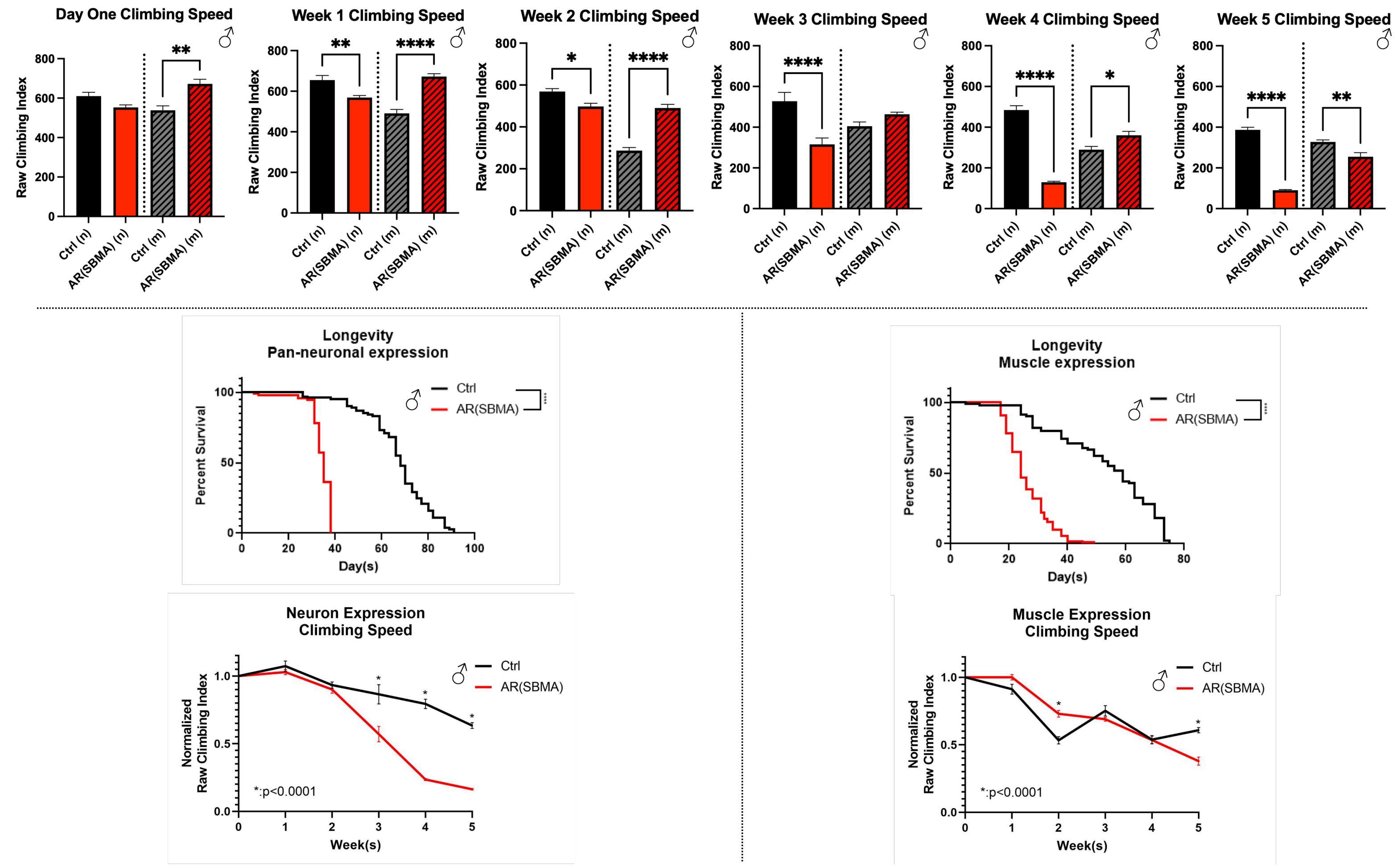

**Supplemental figure 2**

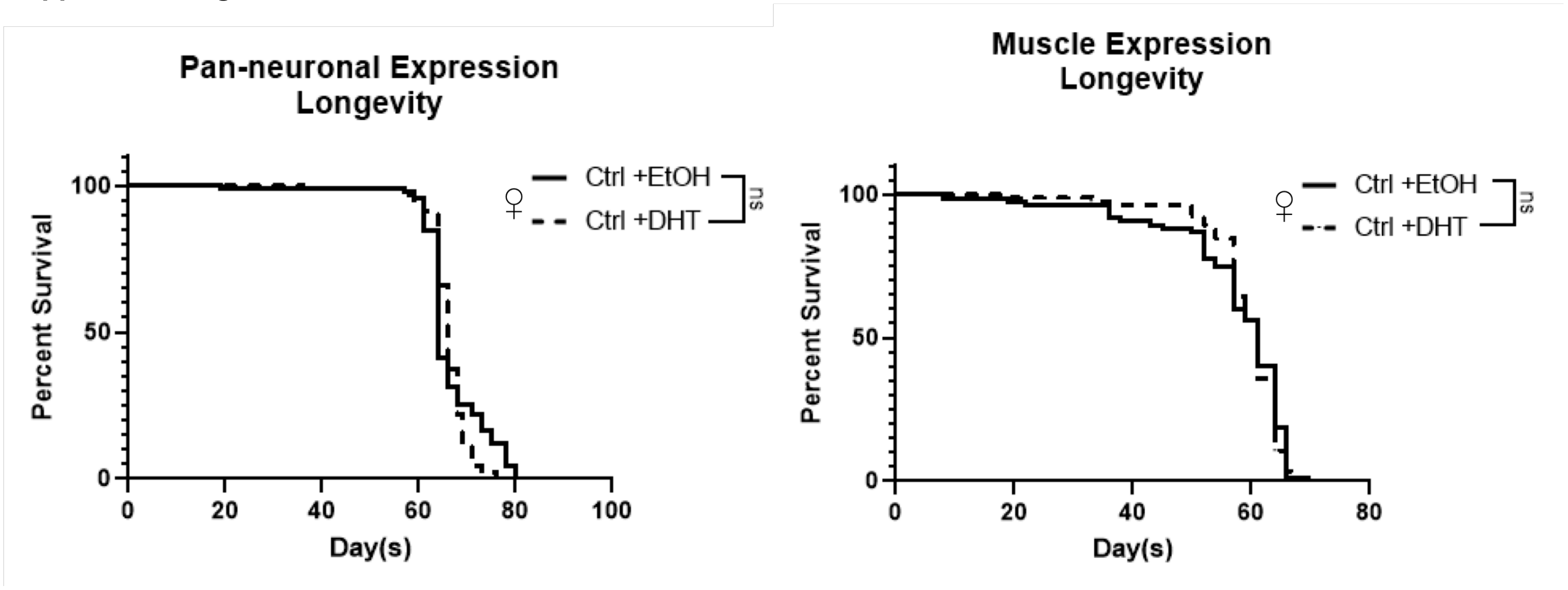

**Supplemental figure 3**

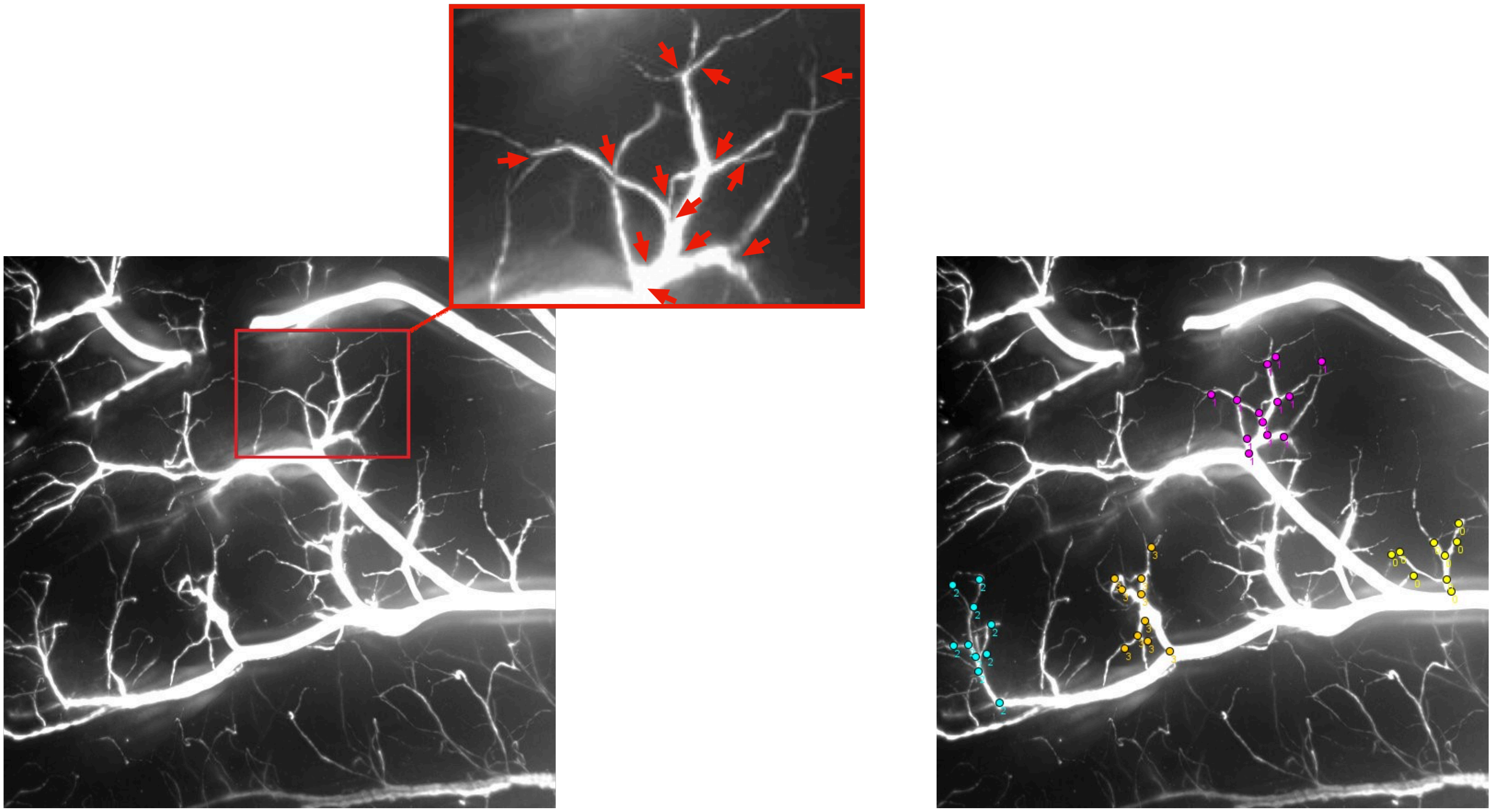

### Supplemental figure 4

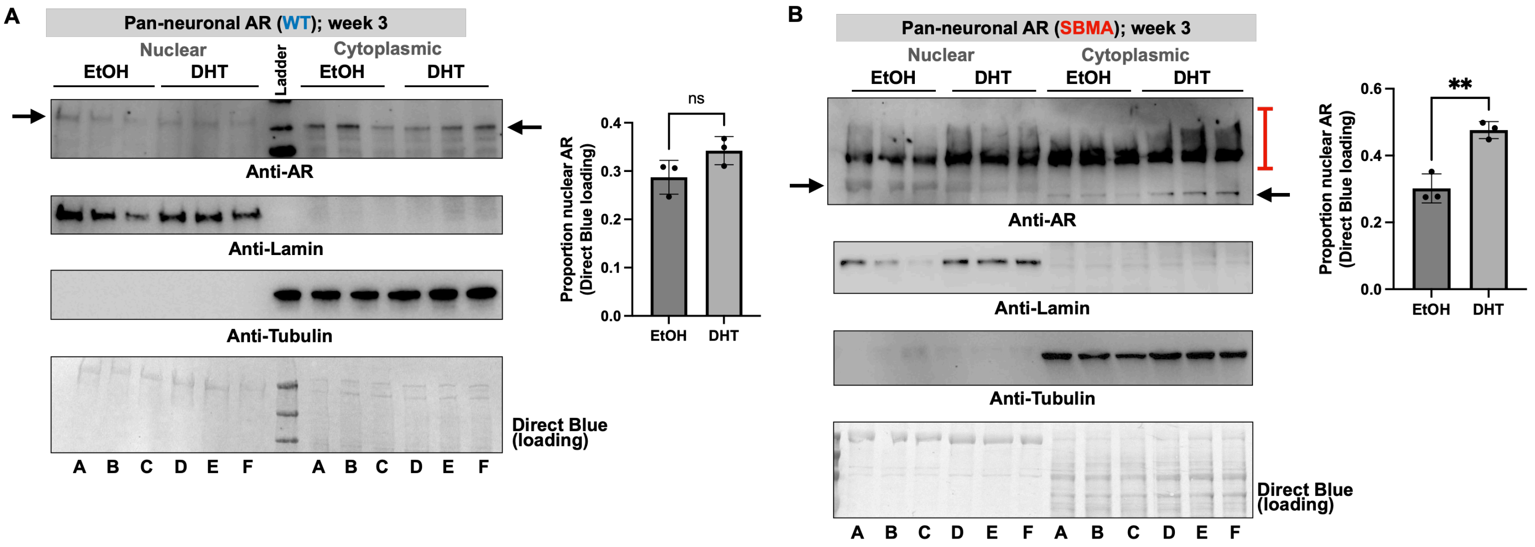
